# A model to study orienting responses in zebrafish, and applications towards the emotion-cognition interaction

**DOI:** 10.1101/827733

**Authors:** Bianca Gomes do Nascimento, Hingrid Suzzan Tarso Oliveira e Oliveira, Hadda Tercya Lima Silva, Diógenes Henrique de Siqueira-Silva, Monica Lima-Maximino, Caio Maximino

**Author notes:** Corresponding author Caio Maximino, Laboratório de Neurociências e Comportamento “Frederico Guilherme Graeff” – Faculdade de Psicologia, Instituto de Estudos em Saúde e Biológicas, Universidade Federal do Sul e Sudeste do Pará, Unidade III, Av. dos Ipês, S/N, s/ CEP, Bairro Cidade Jardim, Marabá/PA, Brazil.

## Abstract

Orienting responses (ORs) are whole-organism reflexes that are elicited by innocuous stimuli, and which decrease in magnitude after stimulus repetition. ORs represent relatively simple responses that can be used to study attentional processes, and are modulated by the organism’s state, including arousal and activation levels, as well as by emotional processes. Here we describe a simple method to study ORs in zebrafish, a model organism increasingly being used in behavioural neuroscience. After presentation of a static visual stimulus, an OR is elicited, characterized by approaching the stimulus and orienting towards it. After repeated stimulation, OR decreases, suggesting habituation. These responses are qualitatively altered by exposure to a fear-eliciting alarm substance (i.e., derived from the skin of a conspecific), since exposed animals avoid the visual stimulus and orient either away from the stimulus or towards it, but at a distance. The protocol can be used to study orienting responses, as well as the impact of fear and arousal on these reflexes.

## 1. Introduction

Zebrafish is an emerging model organism in behavioural neuroscience, and is increasingly being favoured due to its advantages both as a model organism (i.e., ease of upkeep, handling, and reproduction; Key & Devine, 2003) and intermediate physiological and behavioural complexity (Kokel and Peterson 2008; Kalueff et al. 2014; Stewart et al. 2015). As a result, zebrafish has been introduced as a model organism to study the neurochemistry and circuitry of cognition, including executive function and attention (Stewart and Kalueff 2012; Parker et al. 2013). Attention can be described as selective or sustained; the first case refers to the ability to select a target from an array of distractors, while the second is defined as the ability to detect the presence of a target over a prolonged period of time (Parker et al. 2013). Both selective and sustained attention can be measured in different behavioural models, including zebrafish. In the 3-choice serial reaction time task, fish are taught to swim into one of three apertures after maintaining attention to the stimuli (Parker et al. 2012), and is useful to study sustained attention. In the virtual object recognition task (Braida et al. 2014), selective attention is evaluated by observing the time the animal spends exploring novel shapes presented in a screen. In both cases, drugs which increase attention are able to improve performance. Proulx et al. (2014) showed in his work that zebrafish are capable of processing stimuli in parallel search, event though these animals have no telencephalic homologues of the cortical structures which participate in visual search in mammals. This is consistent with the primacy of the superior colliculus/optic tectum for visual search in mammals.

ORs are whole-organism reflexes that are elicited by innocuous stimuli, and which decrease in magnitude (habituate) after stimulus repetition (Pavlov 1927; Sokolov 1963). Orienting responses (ORs) involve a set of behavioural and physiological adjustments that are elicited by novel and/or significant stimuli, leading to the motivational activation of a cascade of perceptual and motor processes that facilitate behavioural selection (Bradley 2009). The virtual object recognition task (Braida et al. 2014) described above is a classical repetition-change paradigm, in which one “truly novel” stimulus is repeatedly presented, and responses to this stimulus are recorded as “orienting responses”, and the decreased responsiveness after a number of presentations is recorded as habituation; at certain intervals, a different stimulus is presented, and orienting responses to this novel stimulus can also be recorded. Interestingly, the optic tectum has been implicated in ORs in goldfish (Torres et al., 2005), suggesting a role in attention and visual search. While ORs are classically interpreted in terms of stimulus novelty, the significance of the stimulus (including its relationship to unconditioned appetitive or aversive cues) is highly important to orienting (Donchin 1981; Barry 2009). The relationship between stimulus significance and ORs is an important factor in judging the attention-emotion interaction.

Attention and emotion can interact through the influence of the first on emotional processing, or through the emotional modulation of the attention processes (Vuilleumier 2005). In that case, once a threat is detected and the initial (automatic) fear response is elicited, further action is needed, demanding the redirection of attentional resources to the threatening stimulus and the engagement of flexible behavioural repertoires. The influence of emotion on spatial attention has been investigated in humans (Beck et al. 1992; Bradley et al. 2000), and non-human animals (e.g., Saiers, Richardson, & Campbell, 1990) and is relevant to understanding psychiatric disorders, given that patients suffering from anxiety disorders present changes in attention and selectivity to threat-associated stimuli (Asmundson et al. 1992; Beck et al. 1992; Bradley et al. 2000; Lautenbacher et al. 2002). Developing methods to study, in non-human animals, attention and ORs that are sensitive to emotional modulation is useful from a translational point of view, increasing the breadth of affected domains in an integrative perspective (Kalueff et al. 2008).

Conspecific alarm substance (CAS) is a complex mixture found in Ostariophysan fishes that is produced by epidermal club cells and released in the water after these cells are injured by, for example, a predator (Maximino et al. 2019). The substance is detected by conspecifics, eliciting defensive (antipredator) behaviour (von Frisch 1938, 1941; Døving and Lastein 2009; Maximino et al. 2019). These behavioural adjustments include erratic swimming and freezing, which increase the probability of detecting a potential threat and decrease the probability of being detected by a predator (Maximino et al. 2019). In that sense, CAS appears to act partially by modulating ORs, either increasing orienting towards novel and/or significant stimuli to promote appraisal, or producing faster responses. In the first case, freezing responses can promote conditions of focused attention towards environmental novelty, increasing detection of the predator; in the second case, fast responses can in fact be detrimental to attention, producing false alarms that are nonetheless counterbalanced by the higher probability of escaping the predator.

The aims of the present article were to 1) propose a simple task to assess ORs in zebrafish towards a novel neutral stimulus (a blue circle); 2) assess whether CAS increases orienting or elicits escape responses away form this novel stimulus. We find that the stimulus induced an approach response, accompanied by orienting towards the stimulus, that habituated after 5 trials. We also find that CAS increased erratic swimming throughout the task, and inverted the direction of movement during stimulus presentation: instead of approaching the stimulus, animals now flew from it. This manuscript is a complete report of all the studies performed to test the effect of a visual stimulus on ORs in zebrafish and of CAS on these orienting responses. We report all data exclusions (if any), all manipulations, and all measures in the study (Simmons et al. 2012).

## 2. Methods

### 2.1. Animals and housing

36 adult (∼4 mo.) mixed sex zebrafish from the longfin phenotype were acquired from local creators and group-housed in tanks with a stock density of 5 animals/2 L. Temperature was kept at 28±2 °C, hardness 75-200 mg/L CaCO_3_, salinity 0.25-0.75 ppt, dissolved oxygen ∼7,8 mg/L at 28 °C, and nitrite levels <0.01 ppm. Animals were kept in the tank for at least two weeks to acclimate before beginning experiments. Environmental characteristics (physico-chemical and health profile) for the housing can be found at GitHub (https://github.com/lanec-unifesspa/lanec-welfare). All manipulations and animal care were in accordance with Brazilian guidelines (Conselho Nacional de Controle de Experimentação Animal - CONCEA 2017)

### 2.2. Sampling plan

Effect sizes for sample size calculation were based on studies on the visual object recognition test (Braida et al. 2014). Since fixed effects (trial and group) were considered more important than the random (subject) effect, sample sizes were calculated based on two-way ANOVAs, with power = 80%, and considering p < 0.05 as a target. After calculation, 12 animals per group were considered sufficient.

### 2.3. Behavioural setup

The experimental setup consisted of two 13 × 13 × 17 (depth x length x depth cm each) glass tanks, positioned side by side, and filled with 2 L tank water. The tanks were positioned at the front of the computer screen (Samsung T20c310lb, 20”, LED screen, nominal brightness 20 cd/m^3^) such that the stimulus appeared in front of the subjects. The computer screen was located at 8.5 cm from the tank centre. Throughout trials, the position of the stimulus was always the same. All sides of the tanks, except the one facing the screen, were covered in opaque white plastic sheets, isolating animals from each other and improving contrast for tracking.

### 2.4. Experimental procedures

A blue circle (R: 0; G: 0; B: 128) was used, based on the ability of zebrafish to discriminate blue from green stimuli in a simple discrimination paradigm (Mueller and Neuhauss 2012). Stimulus size was 72° of horizontal visual angle, calculated considering the diameter of the stimulus in the screen when it is at the from of the animal, and the animal is in the centre of the tank. The size was based on previous work which show that this size is insufficient to produce an escape response in zebrafish larvae (Dunn et al. 2016).

Animals were transferred to the tank and left to acclimate for 3 min before the presentation of the stimuli. The computer screen was turned on for the entire experiment, including acclimatization time. Using an animation based on LibreOffice Impress (v. 6.0.7.3), the stimulus was presented in 1-min intervals (“trials”) interspersed with 1-min stimulus-free periods. In trials in which the stimulus was “on”, the stimulus was presented for the entire duration of the trial. 10 trials were made; total session length was 22 minutes per animal. The number of trials and intertrial durations were chosen based on previous work on arousal in goldfish (Laming and Savage 1980).

### 2.5. Behavioural tracking

A video camera (Sony Handycam® DCR-DVD 610) was positioned above the tanks to allow recording and offline behavioural scoring. All focal fish behaviours were therefore tracked from a top-down perspective, using both TheRealFishTracker (http://www.dgp.toronto.edu/~mccrae/projects/FishTracker/) and fishtracker (https://github.com/joseaccruz/fishtracker). For each behavioural video, a 2D region (arena) was defined for tracking, comprising the inner area of the tank. A region-of-interest (ROI) of 13 x 8.5 cm was determined as “stimulus zone”. The following variables were recorded, using TheRealFishTracker:

- Total time in the stimulus zone (s);
- Absolute turn angle (°), a measure of erratic swimming (Tran et al. 2016);
- Speed (cm/s)

Following Abril-de-Abreu, Cruz, & Oliveira (2015), a region of interest (ROI) of 13 x 3 cm (approximately 25% of the tank), corresponding to the width of the tank and the mean body length of an adult zebrafish, was defined in the area of the arena closer to the computer screen in fishtracker. Animal orientation is defined as a mean projection vector *R*_*proj*_ (Abril-de-Abreu et al. 2015), determined by the transformation of each orientation angle in a vector *r*_*i*_ = (*cos*α_*i*_, *sin*α_*i*_), where α_i_ represents the angle formed by the direction of the centroid-head axis relative to the horizontal axis in each frame. The resulting mean vector, with length *R* = ∥r∥, is a measure of directional focus, and varies between 0 and 1. The projection of this vector *R* in the stimulus direction (arbitrarily defined as 180°) is defined as *R*_*proj*_ = *-Rcosα*. Since *R*_*proj*_ is a mean vector – that is, it takes projection angles in each frame and averages then through the whole 1-min trial –, if the animal is consistently orienting throughout most of the trial, values tend to be positive; if the animal is consistently facing away from the stimulus, values tend to be negative; and if the animal is orienting towards many different directions during the trial, values tend to be close to 0. Thus, positive values indicate direction towards the stimulus, while negative values indicate direction away from the stimulus, and values close to 0 indicate absence of orienting.

During pre-submission peer review, an anonymous referee suggested describing eye use patterns during stimulus presentation. Eye use has been used as a proxy of lateralisation in fish behaviour (Miklósi et al. 1998; Sovrano et al. 1999; Clotfelter and Kuperberg 2007), and has been linked to lateralised brain structures which are involved in fear-like responses (e.g., the habenula; Roussigné, Blader, & Wilson, 2011). Eye use was estimated by analysing projection angles averaged across trials in which the stimulus was “ON” and trials in which the stimulus was “OFF”, with angles between 150° and 179° recorded as “left eye use” (“L”), angles between 181° and 210° recorded as “right eye use” (“R”), and average angles above 210° and below 150° recorded as “no preference”. Thus, eye use was defined as orienting the head in an angle of up to 30°, consistent with what is observed in other experiments on eye use in zebrafish (Miklósi et al. 1998). Since this analysis was added *a posteriori*, and therefore we did not have an *a prior* hypothesis regarding eye use, we relied on descriptive statistics, and therefore results should be considered exploratory.

### 2.6. CAS extraction and exposure

CAS extraction was made according to a protocol described in full detail elsewhere (do Carmo Silva et al. 2018). Briefly, after sacrifice, a single fish was subjected to 15 shallow cuts at both sides of the body, which were subsequently washed with 10 mL distilled water. After removing debris, 7 mL of this solution was separated, and kept on ice until used; this was referred to as “1 unit CAS”. For half of the animals (*n* = 12), CAS was poured directly into the tank water immediately after the animal was transferred to the tank. For the other half (*n* = 12), distilled water was used.

### 2.7. Statistics

For data on time in stimulus zone, absolute turn angle, and speed, a linear mixed-effects model was applied, with subject as random factor, trial as within-subjects factor, and group as between-subjects factor. A temporal autocorrelation structure of order 1 was modelled within levels of the random factor. Analyses were made using the R package ‘nlme’ (v. 3.1-141; https://CRAN.R-project.org/package=nlme). For projection angle data, values were averaged across equal trials (i.e., stimulus “ON” vs. stimulus “OFF”), and a Watson-Williams test for homogeneity of circular data was applied, using the ‘circular’ package (v. 0.4-93; https://r-forge.r-project.org/projects/circular/). For *R*_*proj*_ data, a two-way (treatment X trial type) analysis of variance (ANOVA) was applied, followed by Tukey’s HSD post-hoc test whenever *p* < 0.05. All analyses were run on R 3.6.1 (R Core Team 2019).

### 2.8. Open science practices

Experiments were not formally preregistered. Data and analysis scripts can be found at a GitHub repository (https://github.com/lanec-unifesspa/5-HT-CAS/tree/master/data/behavioral/orienting). Preprint versions of the manuscripts were uploaded to bioRxiv (https://doi.org/10.1101/827733).

## 3. Results

A linear model including subject as random factor was favoured in relation to a model without the random factor (likelihood ratio = 139.5079, p < 0.0001, AIC = 3986.042 vs 4121.55). Main effects of group (χ^2^ = 134.443, p < 2.2 × 10^−16^) and trial (χ^2^ = 64.747, p = 8.52 × 10^−16^), as well as a interaction effect (χ^2^ = 27.905, p = 1.27 × 10^−7^), were found for time in the stimulus zone (Figure 1A). Animals in the control group spend more time near the stimulus when it was “ON”, showing evidence of an orienting response that decreased as the stimulus was repeated; thus, the highest response to the stimulus happened at the beginning of the test, and the response then decreased over repeated expositions.. Animals exposed to CAS, on the other hand, spent *less* time in the stimulus zone when the stimulus was “ON”, suggesting that the stimulus was interpreted as threatening. A main effect of group (F_[1, 524]_ = 305.39, p < 0.0001), but not trial (F_[1, 524]_ = 1.209, p = 0.272), was found for absolute turn angle; no interaction effect was found as well (F_[1, 525]_ = 3.527, p = 0.0717). CAS increased absolute turn angle during the entire test (Figure 1B). No main effects of group (F_[1, 525]_ = 0.019, p = 0.891) or trials (F_[1,525]_ = 0.621, p = 0.431), nor interaction effects (F_[1, 525]_ = 1.2516, p = 0.2638) were found for swimming speed (Figure 1C).

**Figure 1.**
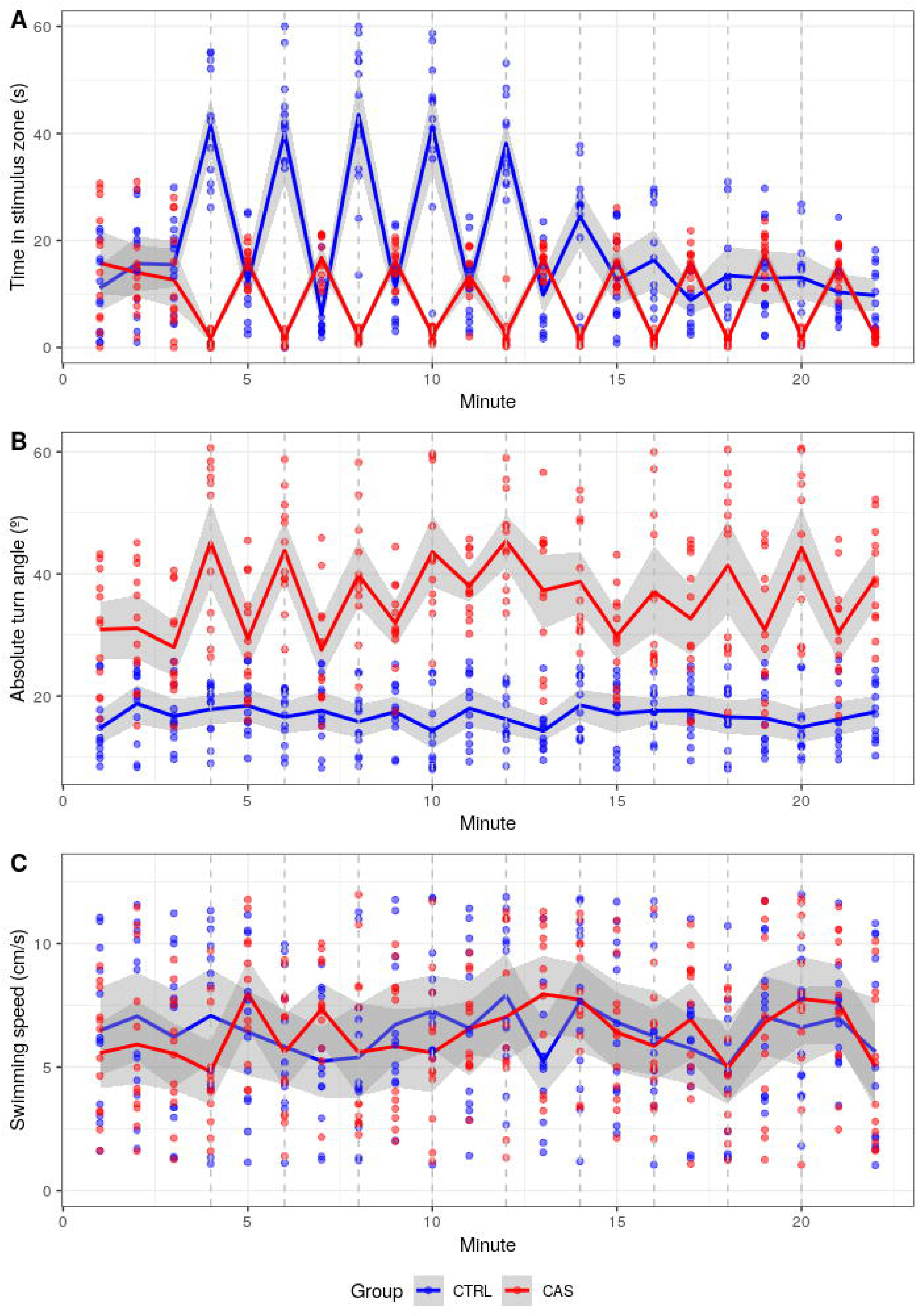
A visual stimulus induces approach responses in zebrafish, while animals exposed to CAS avoid the stimulus and show erratic swimming. Time series scatterplots (*n* = 12/treatment) with (A) time spent near the stimulus, (B) absolute turn angle (erratic swimming), and (C) swimming speed, in 1-min blocks. Stimuli were presented at minutes 4, 6, 8, 10, 12, 14, 16, 18, 20, and 22. Lines and bands represent group averages and bootstrapped 95% confidence intervals around the mean.

For *R*_*proj*_ data, 2 animals from the control group, and three animals from the CAS-exposed group, were removed because the software was not able to detect them. A significant treatment X trial type interaction effect was found for angle (F_[1,36]_ = 13.383, p = 8.07×10^−4^; Figure 2A). A significant interaction was also found for *R*_*proj*_ (F_[1, 34]_ = 88.639, *p* = 5.34×10^−11^; Figure 2B); in the control group, the presentation of the stimulus increased *R*_*proj*_ (*p* < 0.0001), while in the CAS group stimulus presentation decreased *R*_*proj*_ (*p* = 0.000006).

**Figure 2.**
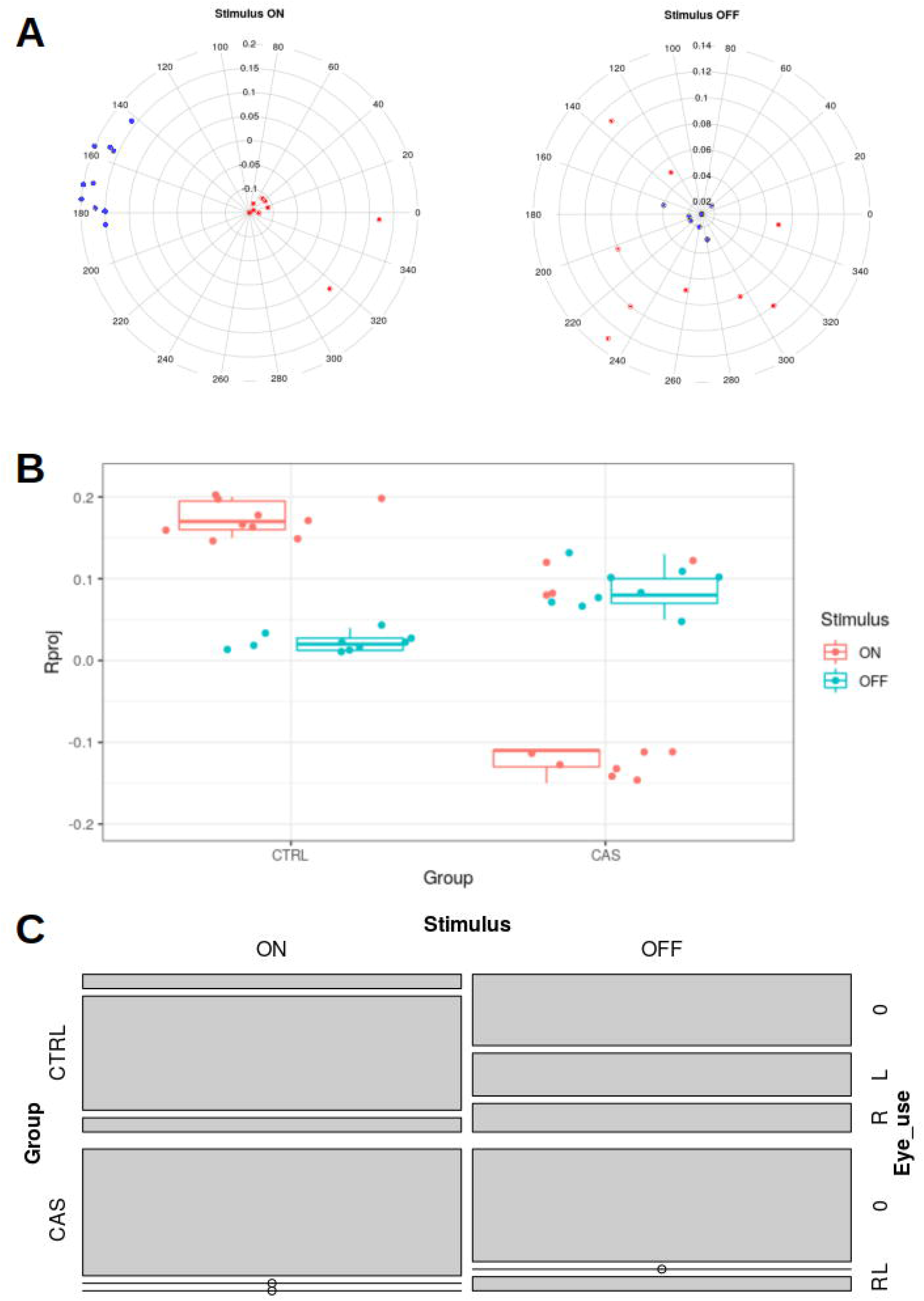
Orienting responses towards a visual stimulus in zebrafish, exposed or not to an alarm substance. (A) Polar plots of the focal fishes’ individual mean resultant vector’s angles (0° to 360°, stimulus positioned at 180°), combined with corresponding vector lengths *R* (0 to 1) for each treatment (control – black dots; CAS – red dots); when the stimulus is on, animals from the control group significantly deviate from a uniform distribution, clustering around its group mean resultant vector. (B) Scatter plots (*n* = 9-12/treatment) of the individual (coloured dots) resultant vector’s lengths *R* projected onto the stimulus direction (*R*_*proj*_), and corresponding group summaries (boxplots), for each treatment; positive values indicate directional focus towards the stimulus, zero indicates no directionality, and negative values indicate focus opposite to the stimulus. (C) Mosaic plots (*n* = 9-12/treatment) of the proportions of animals observing the stimulus with the left eye (“L”), right eye (“R”), or no preference (“0”), across trial types (stimulus ON vs. stimulus OFF) and groups (control vs. CAS).

Finally, in animals from the control group, a preference for left eye use was observed in trials in which the stimulus was turned on, while no preference was apparent in trials in which the stimulus was turned off (Figure 2C). During CAS exposure, no preference was apparent in any trial type.

## 4. Discussion

In this work we demonstrate a simple method to elicit orienting responses in zebrafish towards a visual stimulus. The orienting response was characterized by an approach to the stimulus, as well as body orientation towards the stimulus when it was presented. These orienting responses follow the classical pattern of habituation after repeated stimulus presentation, suggesting that they accurately capture orienting. Moreover, we show that high increases in arousal by exposing animals to an alarm substance abolish the orientation response and substitute it for a defence response.

Orienting responses are relatively simple and amenable to manipulations of different components, and have translational value to understand the emotion-cognition interaction (Bradley 2009). Orienting responses allow a preliminary processing of the environment in terms of stimulus significance, and selectively filter irrelevant information that arrives through sensory channels (Sokolov 1963; Barry 2009). During stimulus presentation, zebrafish in the control group responded with an orienting response, approaching the stimulus and changing body orientation to face it. As stimulus presentation continued, this response showed signs of habituation. Thus, in the initial trials, orienting responses allowed sensory information to be analysed by perceptual areas, which results in information about the stimulus being available to perceptual processes. As the novelty of the stimulus decreases, habituation processes lead to a decline in the orienting response (Pavlov 1927; Sokolov 1963). The decrease in responding is likely not due to increased difficulty in discriminating stimuli, as approach times do not decrease with increasing number of distractors in a parallel visual search task (Proulx et al., 2014).

Orienting responses are in part mediated by the locus coeruleus, and noradrenaline release in the forebrain will facilitate sensory processing, enhance cognitive flexibility and executive function, and promote memory consolidation in limbic structures (Sara and Bouret 2012). Since the ability to attend to discrete pieces of information is limited, due to the high density of stimuli available to the visual system, ORs direct attention to certain areas of the visual field, in the absence of eye movements (Wright & Ward, 1998).

Stednitz et al. (2018) used orienting responses towards conspecifics to dissect mechanisms of social attention in zebrafish, and found that lesions in cholinergic neurons of the ventromedial telencephalon decreased the orienting response towards conspecifics. While it is not known whether these cholinergic neurons also participate non-social orienting responses, acetylcholine and noradrenaline both have been implicated in the effects of arousal in attention. Alternatively, the optic tectum – which has been shown in goldfish to participate in both orienting and escape responses (Torres et al., 2005) – can be a site of convergence for these mechanisms.

Arousal levels are thought to amplify the phasic orienting response by sensitizing the organism to both significant and non-significant stimuli (Barry 2009). In the present work, however, orienting was not simply amplified by CAS, but qualitatively changed it. During stimulus presentation, animals avoided it, spending more time away from the stimulus and either orienting away from it or towards it. Moreover, this response did not habituate throughout trials, and was observed also in the absence of the stimulus – although it was significantly increased by stimulus presentation. CAS also elicited an increase in absolute turn angle, suggesting erratic swimming, as observed in zebrafish exposed to CAS and other fear-eliciting stimuli (Gerlai et al. 2009; Ahmed et al. 2012; Maximino et al. 2019). CAS is known to elicit fear-like behaviour in fish from the Ostariophysan superorder, decreasing the probability of predation by promoting escape or freezing, and increasing the probability that conspecifics detect a predator by inducing increased arousal and attention (Smith 1992; Maximino et al. 2019). In rats, increasing arousal by placement in a novel environment (Richardson et al. 1988) or administering electric shocks (Saiers et al. 1990) disrupts the orienting response, suggesting either competing mechanisms or the induction of an emotional state. These results suggest that CAS-induced arousal did not merely amplify the orienting response, but induced a state of hypervigilance, in which a non-significant stimulus is interpreted as threatening. To understand whether CAS elicits competing responses (i.e., “sensory overloading”) or an emotional state, using other olfactory stimuli which induce attraction responses (e.g., bile salts; Krishnan et al., 2014) would be interesting, but that falls outside the scope of this work.

The present results also suggest lateralisation of orienting responses. In animals in the control group, orienting responses were biased towards left-eye use, an effect which was not observed when the stimulus was off. However, since animals in the group that was exposed to CAS generally escaped the stimulus, it was not possible to infer whether lateralisation was present in these responses. Zebrafish have been shown to first inspect a novel stimulus using the right eye, and then using left frontal viewing as the object is further presented (i.e., as novelty decreases)(Miklósi et al. 1998). While preliminary and exploratory, our results suggest that orienting responses are lateralised, which could implicate brain areas known to be involved in behavioural lateralisation (e.g., the habenula-interpeduncular nucleus system; Concha, 2004) in orienting in zebrafish.

Taken in conjunction, our results suggest a protocol to evoke orienting responses in zebrafish as a method to study attention, as well as an activating effect of the alarm substance in these responses. The results underline an important emotion-cognition interface in zebrafish that can be exploited to study behaviour and that has implications to using zebrafish behavioural models to study biological psychiatry. Future work is needed to understand the relationship between these effects and arousal-associated molecules.

## Acknowledgements

This work was supported by a Conselho Nacional de Desenvolvimento Científico e Tecnológico (CNPq/Brazil) grant (Edital Universal 2016, #400726/2016-6). BGN was the recipient of a CNPq/Brazil scholarship.

